# Bridging the Gap: biofilm-mediated establishment of *Bacillus velezensis* on *Trichoderma guizhouense* mycelia

**DOI:** 10.1101/2024.06.06.597722

**Authors:** Jiyu Xie, Xinli Sun, Yanwei Xia, Lili Tao, Taimeng Tan, Nan Zhang, Weibing Xun, Ruifu Zhang, Ákos T. Kovács, Zhihui Xu, Qirong Shen

## Abstract

Bacterial-fungal interactions (BFIs) are important in ecosystem dynamics, especially within the soil rhizosphere. The bacterium *Bacillus velezensis* SQR9 and the fungus *Trichoderma guizhouense* NJAU 4742 have garnered considerable attention due to their roles in promoting plant growth and protecting their host against pathogens. In this study, we utilized these two model microorganisms to investigate BFI. We firstly demonstrate that while co-inoculation of *B. velezensis* and *T. guizhouense* could promote tomato growth, these two microorganisms display mutual antagonism on agar solidified medium. To resolve this contradiction, we developed an inoculation method, that allows *B. velezensis* colonization of *T. guizhouense* hyphae and performed a transcriptome analysis. During colonization of the fungal hyphae, *B. velezensis* SQR9 upregulates expression of biofilm related genes (e.g. *eps, tasA*, and *bslA)* that is distinct from free-living cells. This result suggested an intricate association between extracellular matrix expression and hyphae colonization. In accordance, deletion *epsD, tasA, or* both *epsD* and *tasA* genes of *B. velezensis* diminished colonization of the *T. guizhouense* hyphae. The insights from our study demonstrate that soil BFIs are more complex than we understood, potentially involving both competition and cooperation. These intricate biofilm-mediated BFI dynamics might contribute to the remarkable diversity observed within soil microbiota, providing a fresh perspective for further exploration of BFIs in the plant rhizosphere.

## 1 Introduction

In nearly all ecosystems, bacteria and fungi frequently assemble into dynamic co-evolving communities, encompassing microbial species from a wide diversity of fungal and bacterial families [1]. Bacterial-fungal interactions (BFIs) are vital in ecosystems, driving biochemical cycles and impacting the health of plants and animals [2,3]. The resulting by-products have diverse applications in agriculture, forestry, environmental protection, food processing, biotechnology, and medicine, offering solutions to various challenges and fostering innovation [4].

The pattern of bacterial and fungal abundances arise from intimate biophysical and metabolic interactions, leading mutual dependency and co-evolution between these microbial partners [5]. In soil, *Pseudomonas fluorescens* BBc6R8 enhances the survival of the mycorrhizal fungus *Laccaria bicolor* in challenging soil conditions [6], and *L. bicolor* also supports the survival of the bacterium [7]. BFIs can facilitate the colonization of surfaces that would otherwise be inaccessible to certain microorganisms. Using an *in vitro* polystyrene-serum system, *Candida albicans* was shown to strongly enhance biofilm formation of *Staphylococcus aureus*, where the bacterial cells associated with the fungal hyphae rather than the plastic substrate[8].

*Trichoderma* and *Bacillus* as the most extensively studied PGPMs (plant growth promoting microorganisms) and these have been extensively studied to elucidate the mechanisms for plant growth promotion [9,10]. *Trichoderma guizhouense*, an opportunistic and non-pathogenic plant symbiotic fungus, directly inhibits plant pathogens through parasitism [11] and indirectly by inducing local and systemic defenses in plants [12], which ultimately lead to enhanced root development and plant growth. *Bacillus velezensis*, a Gram-positive soil-dwelling bacterium, is non-pathogenic and commonly resides in association with the plant rhizosphere [13]. To exert its beneficial effects, *Bacilli* rely on their ability to form biofilm. Several studies reported that the co-inoculation of *Trichoderma* sp. and *Bacillus* sp. into the rhizosphere enhance their plant-growth-promoting abilities and disease suppression [14,15]; however, the underlying mechanisms have not been thoroughly investigated.

Biofilms are the collective lifestyle of most microorganisms on earth, where cells are embedded within a self-produced extracellular polymeric substances adhering to each other and surface [16]. The matrix contributes to the structure of a biofilm, allowing for the interaction of multiple organisms in close proximity, including the exchange of metabolites, signaling molecules, genetic material, and defensive compounds [17,18]. However, research on BFIs within biofilms is currently limited due to significant differences in growth rates, patterns, and other factors between different bacteria and fungi.

In this study, we describe that the co-inoculation of *T. guizhouense* NJAU 4742 and *B. velezensis* SQR9 exhibit a synergistic growth-promotion of tomato plants. To elucidate this interaction, we established a cultivation method which designed to mitigate the impact of growth rate disparities. Transcriptome analysis revealed that hyphae-colonized *B. velezensis* display a distinct gene expression profile, including biofilm formation related genes being upregulated. Through construction of specific biofilm mutants, stereo microscopy, confocal laser scanning microscopy (CLSM) and scanning electron microscopy (SEM), we describe that biofilm formation by *B. velezensis* is essential for colonization on *T. guizhouense* hyphae.

## 2 Material and methods

### 2.1 Strains and growth conditions

The bacterial and fungal strains in this study are listed in Table 1. *B. velezensis* strain SQR9 (NCBI accession PRJNA227504) were grown at 37 °C in Lysogeny Broth medium (LB-Lennox, Carl Roth, Germany; tryptone10 g/L, yeast extract 5 g/L and NaCl 5 g/L) supplemented with 1.5% Bacto agar if required. *T. guizhouense* strain NJAU 4742 (NCBI accession PRJNA314460) were grown at 28 °C in Potato Dextrose Agar (PDA, BD Difco, USA) medium for 7 days with light. Spores were harvested with 5 ml water and filtered through Miracloth. Spores were stored at 4 °C until use. When required, antibiotics were added to media at the following final concentrations: chloramphenicol (Cm) at 5 µg/mL, erythromycin (Em) at 1 µg/mL, spectinomycin (Spec) at 100 µg/mL and Hygromycin B (Hyg) at 100 µg/mL.

**Table 1.**
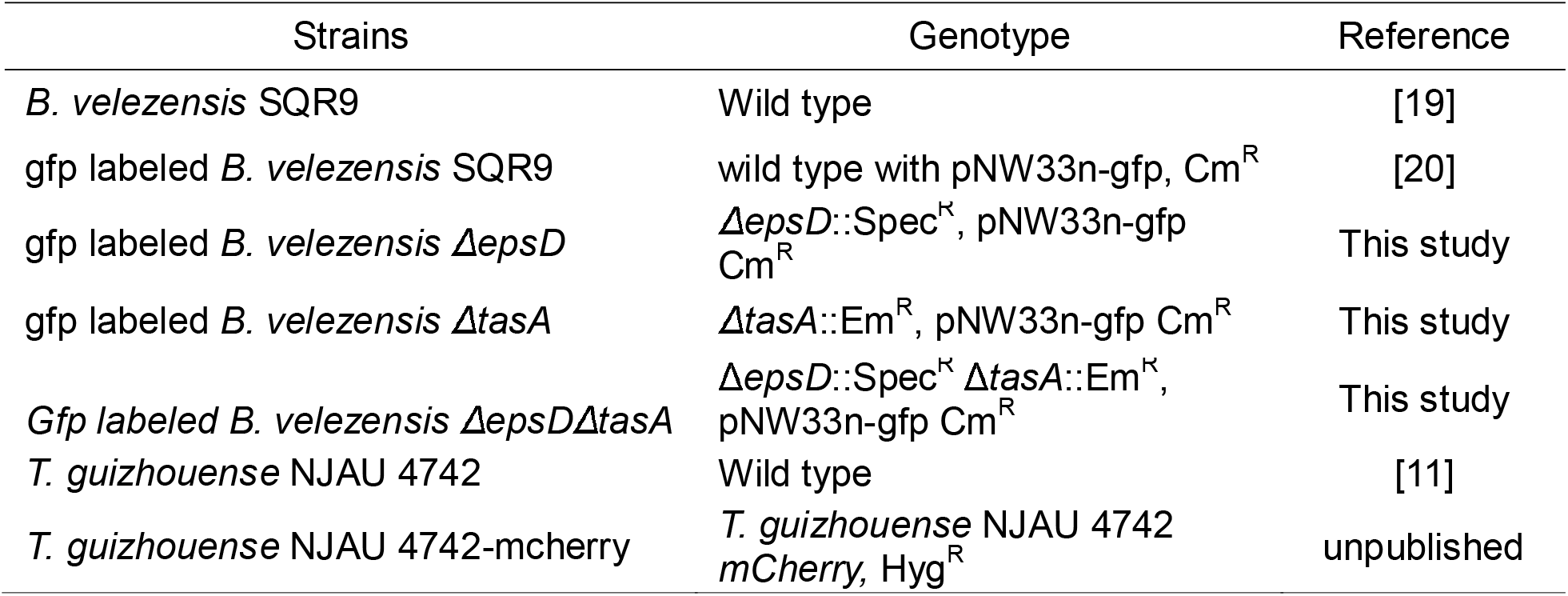
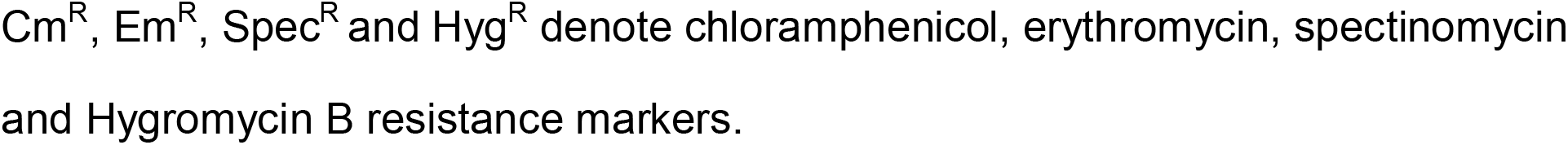
Bacterial and fungal strains used in this study.

| Strains | Genotype | Reference |
| --- | --- | --- |
| *B. velezensis* SQR9 | Wild type | [19] |
| gfp labeled *B. velezensis* SQR9 | wild type with pNW33n-gfp, Cm<sup>R</sup> | [20] |
| gfp labeled *B. velezensis* $\Delta epsD$ | $\Delta epsD::Spec^K$, pNW33n-gfp<br>Cm<sup>R</sup> | This study |
| gfp labeled *B. velezensis* $\Delta tasA$ | $\Delta tasA::Em^R$, pNW33n-gfp Cm<sup>R</sup> | This study |
| Gfp labeled *B. velezensis* $\Delta epsD\Delta tasA$ | $\Delta epsD::Spec^K$ $\Delta tasA::Em^K$,<br>pNW33n-gfp Cm<sup>R</sup> | This study |
| *T. guizhouense* NJAU 4742 | Wild type | [11] |
| *T. guizhouense* NJAU 4742-mcherry | *T. guizhouense* NJAU 4742<br>*mCherry*, Hyg<sup>R</sup> | unpublished |

Cm<sup>R</sup>, Em<sup>R</sup>, Spec<sup>R</sup> and Hyg<sup>R</sup> denote chloramphenicol, erythromycin, spectinomycin and Hygromycin B resistance markers.

### 2.2 Bacteria-fungi coculture assay

4 µL spores (10^8^ spores/mL) of *T. guizhouense* were inoculated into a single well of 6 well plates with 4 ml LB medium and incubated at 28 °C with shaking at 180 rpm. After 24h, 4 µL *B. velezensis* culture with an optical density of 1.0 at 600 nm (OD_600_=1.0) was inoculated into each well. The plates were incubated at 28 °C for 24h with shaking at 180 rpm. Following incubation, the fungal pellets were collected (Fig 1A) and washed three times with distilled water. The samples were used for Microscopy observation or RNA extraction.

**Fig 1.**
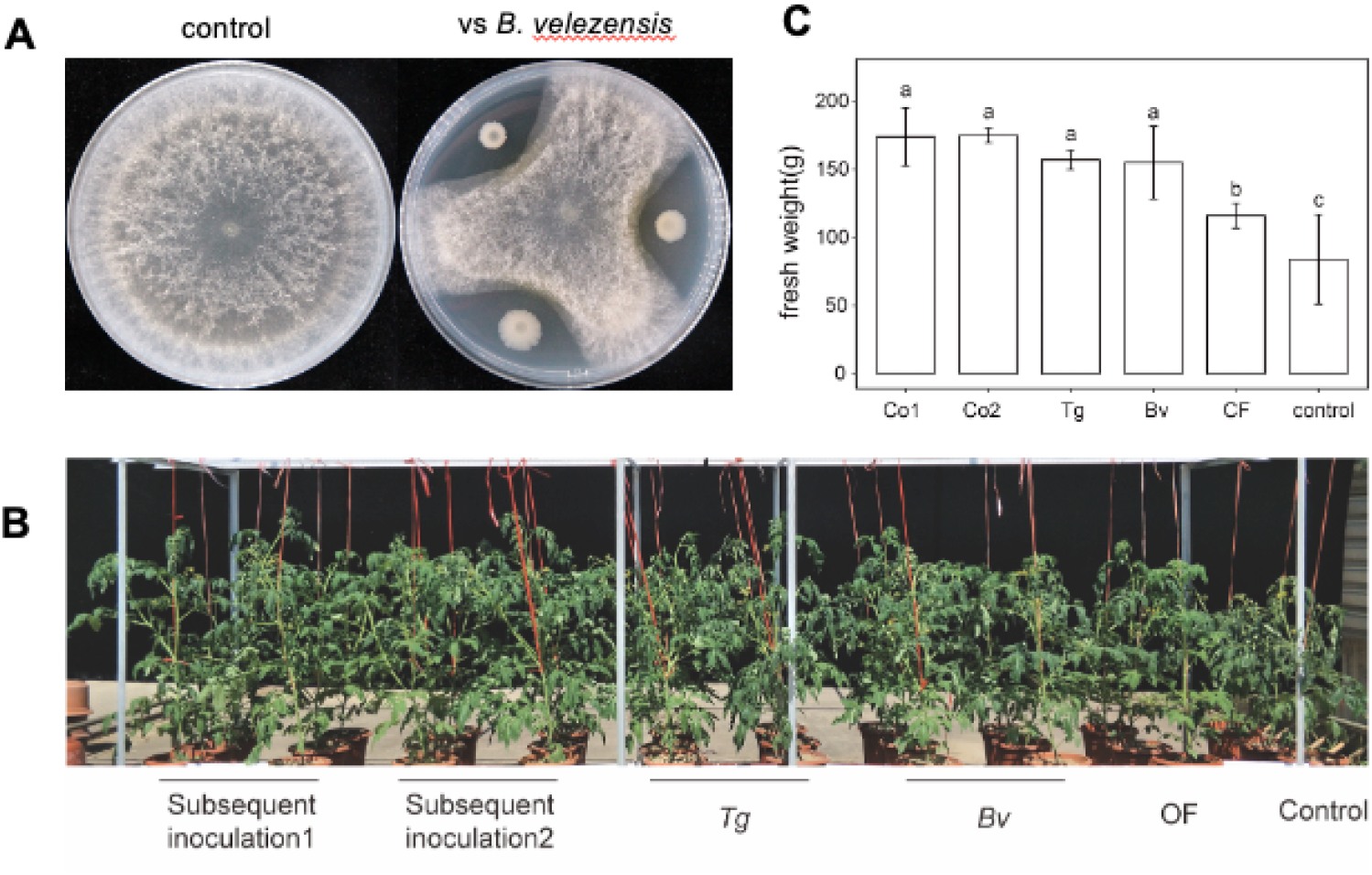
Enhanced growth of tomato under the coculture of *B. velezensis* and *T. guizhouense*. **(A)** The interaction between *T. guizhouense* and *B. velezensis* on agar plates. The diameter of petri dish plate: 9 cm. **(B)** Fresh weight of Plants. **(C)** Tomato plants with different treatments. Control: water; CF: chemical fertilizer; Bv: *B. velezensis*; Tg: *T. guizhouense*; Subsequent inoculation1(Co1): inoculated *B. velezensis* in the soil first then *T. guizhouense*; Subsequent inoculation1 2(Co2): inoculated *T. guizhouense* in the soil first then *B. velezensis*. Bars represent the mean ± s.d. (n=3). Significance test was performed using two-way ANOVA followed by Tukey’s post hoc test via Prism 10. Different letters indicate statistically significant (p < 0.05) differences.

### 2.3 RNA-seq analysis

Total RNA was extracted using the E.Z.N.A. bacterial RNA kit (Omega Bio-tek, Inc.). RNA integrity was assessed using the RNA Nano 6000 Assay Kit of the Bioanalyzer 2100 system (Agilent Technologies, CA, USA), and then sequenced on TruSeq PE Cluster Kit v3-cBot-HS (Illumina) platform. Raw sequencing data have been deposited to the NCBI SRA database under BioProject accession number PRJNA1111338. Raw reads were quality-trimmed and then mapped to reference genomes using Bowtie2-2.2.3 software[21]. HTSeq v0.6.1 was used to count the reads numbers mapped to each gene. And then FPKM of each gene was calculated based on the length of the gene and reads count mapped to this gene. Differential expression analysis was performed using the DESeq2 R package[22] The resulting p values of genes were adjusted using Benjamini and Hochberg’s approach for controlling the false discovery rate (FDR). Genes were assigned as differentially expressed when log2 fold change (LFC) > 2 and FDR < 0.05. For functional analysis, the protein-coding sequences were mapped with KEGG Orthology terms using EggNOG-mapper v2[23]. P values of pathways were corrected for multiple hypothesis testing using the Benjamini and Hochberg’s approach.

### 2.4 Microscopy

A confocal laser scanning microscope Axio Observer Z1/7 (Carl Zeiss) equipped with a 20×/0.50 M27 EC Plan-Neofluar objective, was used to acquire confocal microscopic images. *B. velezensis* was excited with 488nm laser and detected at 509nm for GFP signal. A stereo-microscope Axio Zoom V17 (Carl Zeiss) equipped with a 1.0× Plan-Neofluar objective, was used to acquire images of plates (with different magnifications). Fluorescence of *B. velezensi*s and *T. guizhouense* samples were detected at 488nm(Ex) and 409nm (Em) and 587nm (Ex) and 610nm (Em) for the GFP and mCherry signals, respectively. Images were collected with Zen software (Carl Zeiss) and analyzed using the ImageJ software. For scanning electron microscopy, the samples were fixed with 2.5% electron microscopy grade glutaraldehyde and 2% osmium tetroxide (OsO) in distilled water. The sample were dried using tert-butanol and ethanol. After coating, the samples were imaged with a Scanning Electron Microscope Regulus 8100 (Hitachi High-Tech, Tokyo, Japan).

### 2.5 Construction of gene deletion bacterial strains

Deletion of *tasA* and *epsD* genes was performed using overlap-PCR based strategy as previously described [24]. The upstream (800 bp) and downstream regions (1000 bp) were amplified from wild-type *B. velezensis* genome using primer pairs (Table S1) tasA_UF/tasA_UR, epsD_UF/eps_UR and tasA_DF/tasA_DR epsD_DF/epsD_DR, respectively. The antibiotic markers (Em and Spec) were amplified from plasmids pPax01[25] and pheS-SPC[26] using the primer pairs Em_F/Em_R and Spc_F/Spc_R, respectively. The three fragments were fused using overlay PCR in the order of upstream, antibiotic region and downstream fragments. The resulting *tasA* and *epsD* deletion amplicons (PCR products) were directly transformed into *B. velezensis* containing the constitutive GFP (*B. velezensis* SQR9-gfp), and the transformants were selected on LB agar medium containing erythromycin, spectinomycin, or both erythromycin and spectinomycin, to obtain the *tasA, epsD* or *tasA-epsD* deletions, respectively. Mutants were further confirmed by DNA sequencing.

## 3 Results

### 3.1 Co-inoculation of *B. velezensis* and *T. guizhouense* maintains their plant growth-promotion

In laboratory conditions, co-incubation of *B. velezensis* and *T. guizhouense* on PDA agar medium revealed a strong competitive interaction (Fig 1A), attributed to the antifungal secondary metabolites produced by *B. velezensis* [27]. Such inhibitory activity between the bacterium and the fungus questions whether the two species can co-exist when applied as plant growth promoting agents. Subsequent inoculation of these two PGPM might reduce inhibition between them.

To investigate the effect of subsequent co-inoculation of *B. velezensis* and *T. guizhouense* on tomato growth, we used a pot experimental setup as depicted in Fig S1. In Subsequent-inculation1, we initially incubated two-week-old tomato plant seedlings with *B. velezensis* and incubated for one week, followed by subsequent inoculation of *T. guizhouense*. Conversely, Subsequent-inculation2 involved the reverse experiments. After six weeks of cultivation, notable differences in tomato growth were recorded (Fig 1B). The visual plant growth as well as the plant fresh weight quantification demonstrated that treatments with PGPM significantly enhanced plant growth compared with the control and the chemical fertilizer treatments. Interestingly, despite mutual antagonism, *B. velezensis* and *T. guizhouense* were able to enhance tomato growth when co-inoculated to the roots of tomato plants.

### 3.2 *B. velezensis* can colonize the pre-grown hyphae of *T. guizhouense in vitro*

To elucidate the contradiction between the antagonism observed between *B. velezensis* and *T. guizhouense* on agar medium and their ability to facilitate plant growth when inoculated subsequently, we initiated an in-depth investigation into their interactions under laboratory settings. Considering the growth rate of fungi and bacteria might differ on laboratory media, we hypothesized that these two strains could achieve co-existence through controlling the time of inoculation. Here, the slower growth rate of the fungus, *T. guizhouense* might carry a disadvantage during this specific BFI. Thus, we implemented a cultivation method [28,29] aiming at alleviating the competitive pressure resulting from tentative differential growth rates (Fig 2A). Initially, we inoculated the spores of mCherry-labeled *T. guizhouense* in a 6-well plate, allowing for spore germination during a 24-hour period with shaking. After a day of incubation at 28 °C, 20-30 hyphal pellets of *T. guizhouense* were observed in each well. Subsequently, GFP-labeled *B. velezensis* was directly inoculated into the fungal cultures and the co-culture was incubated for 24 hours.

**Fig 2.**
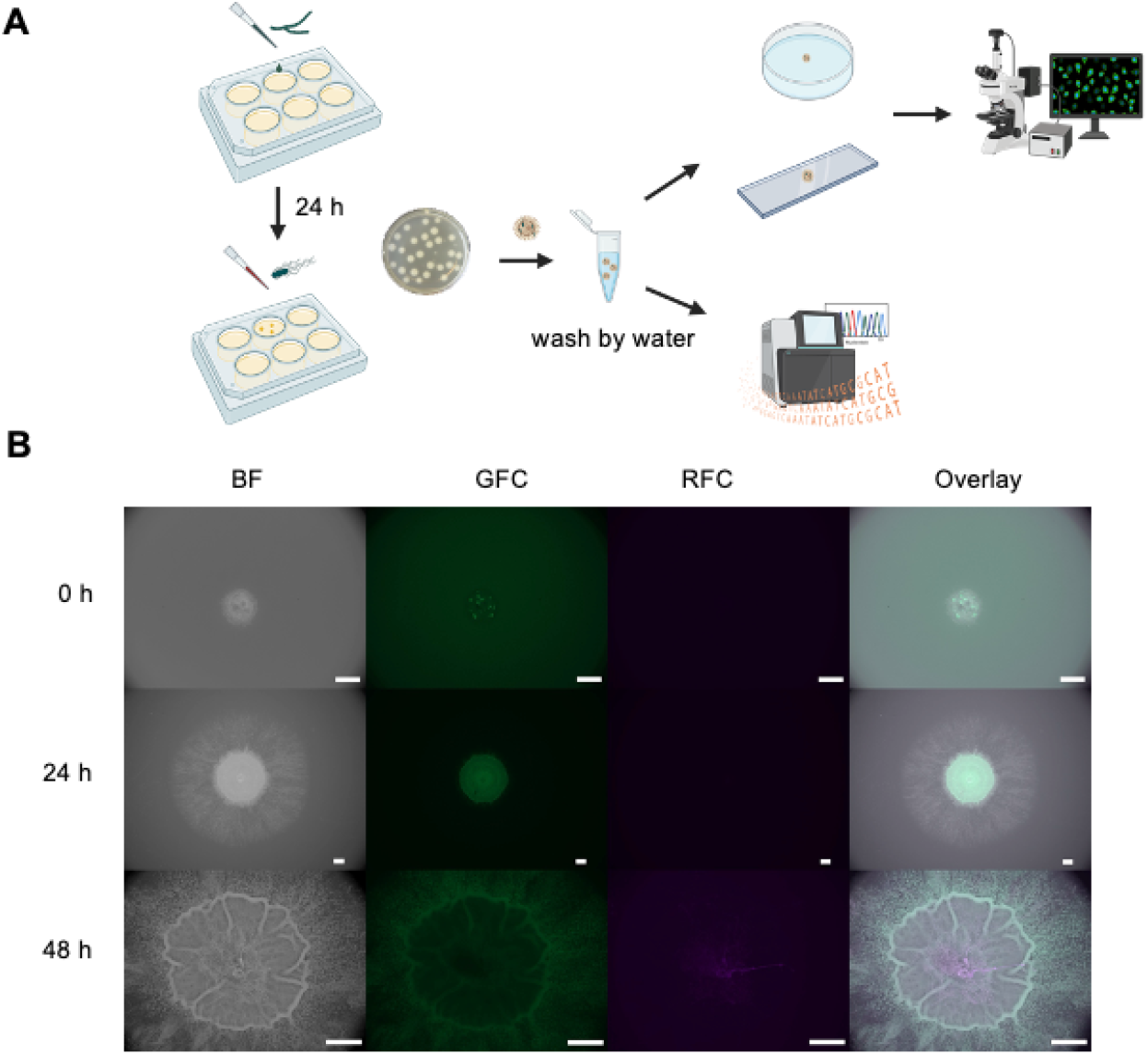
Colonization of the *T. guizhouense* hyphae by *B. velezensis*. **(A)** The experiments setup and analysis method. **(B)** Co-culture images of *B* .*velezensis* (gfp) and *T. guizhouense* (mCherry) by CLSM. Images were taken in 0h, 24h, 48h. BF: bright channel; GFC: green fluorescent channel; RFC: red fluorescent channel. Scale bars indicate 1000 μm.

To dissect whether this preincubation step facilitates interaction among the two PGPM, the non-attached *B. velezensis* cells were first removed by washing the hyphae pellets using sterile water and BFI pellet was placed onto PDA medium. Surprisingly, we discovered that this approach allowed the simultaneous growth of both *B. velezensis* and *T. guizhouense* (Fig 2B). Although only a small amount of green fluorescence was observed at the start, after 24 hours of incubation, the presence of *B. velezensis* was evident in the center of the colony, while *T. guizhouense* spread radially outwards. This observation suggests that within such coculture setup, *T. guizhouense* and *B. velezensis* lack direct inhibition of each other. After 48 hours of incubation, *B. velezensis* formed a biofilm at the center whilst the hyphae of *T. guizhouense* further extended towards the non-colonized region of the agar medium. In summary, when *T. guizhouense* was pre-cultivated before co-cultivation with *B. velezensis*, an interaction could be observed that enabled *B. velezensis* to colonize the hyphae of *T. guizhouense* and continue to grow in subsequent cultivations.

### 3.3 *B. velezensis* differentially regulates biofilm and motility-related genes during colonization of the fungal hyphae

To gain a detailed insights into the hyphae colonization mechanism by the bacterium, the transcriptome of *B*. velezensis was compared using three conditions: control (bacterial monoculture), free-living bacterial cells in co-culture, and hyphae attached bacterial cells (Fig 3A). First, principal coordinates analysis (PCoA) was employed to visualize the transcriptome differences of the bacterial cultures under these conditions using Bray-Curtis distances (Fig 3B). The three sample types were clearly separated on the coordinates. Compared with the monoculture samples, the hyphae-colonized *B. velezensis* cells exhibited downregulation of genes from 12 functional categories and upregulation of genes from 8 functional categories (Fig 3C). Major downregulated functional categories included ribosome synthesis, flagellar assembly, bacterial chemotaxis, and histidine metabolism. The decreased expression of ribosome and histidine metabolism suggests a reduced rate of protein synthesis and cellular growth [30]. Furthermore, the reduced expression of bacterial genes encoding flagellar assembly [31] and bacterial chemotaxis [32] indicates a decreased cell motility, consistent with the sessile colonization of the fungal hyphae. The major upregulated functional categories included phosphotransferase system, indicating that cells were adapting to the new environment by adjusting energy metabolism pathways. Additionally, comparison of free-living cells in the coculture and the monoculture, 11 up and 8 downregulated functional categories could be detected in the cocultured *B. velezensis*. The category with the highest number of differential gene expression included the biosynthesis of secondary metabolites [33], which suggests a bacterial stress response to inhibit the fungus (Fig S3).

**Fig 3.**
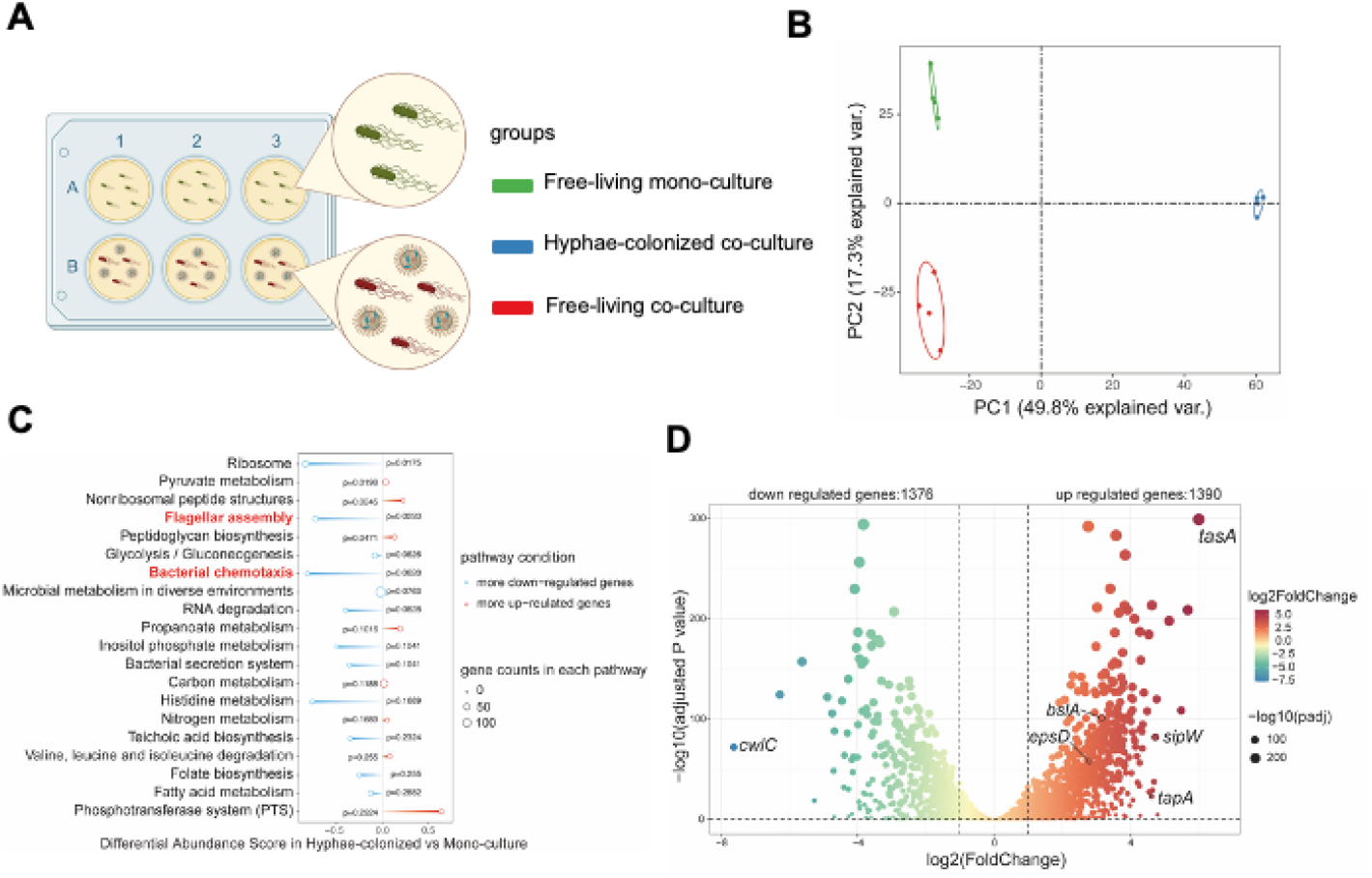
Transcriptome analysis of *B. velezensis*. **(A)** Schematic representation of transcriptome treatments. **(B)** Principal Coordinates Analysis (PCoA) of *B. velezensis* is plotted based on the Bray−Curtis distance metrices for taxonomical data (p < 0.01). Permutational multivariate analysis of variance (PERMANOVA) was performed using the adonis function from the vegan R package. Samples were collected form different treatment (n=4).**(C)** KEGG enriched pathway analysis of hyphae-colonized *B. velezensis* compared with mono-culture(control). Blue indicated there were more down-regulated genes in the pathway, red indicated there were more up-regulated genes in the pathway. **(D)** The log2(fold change) of genes in the colonized *B. velezensis* compared with mono-culture.

**Fig 4.**
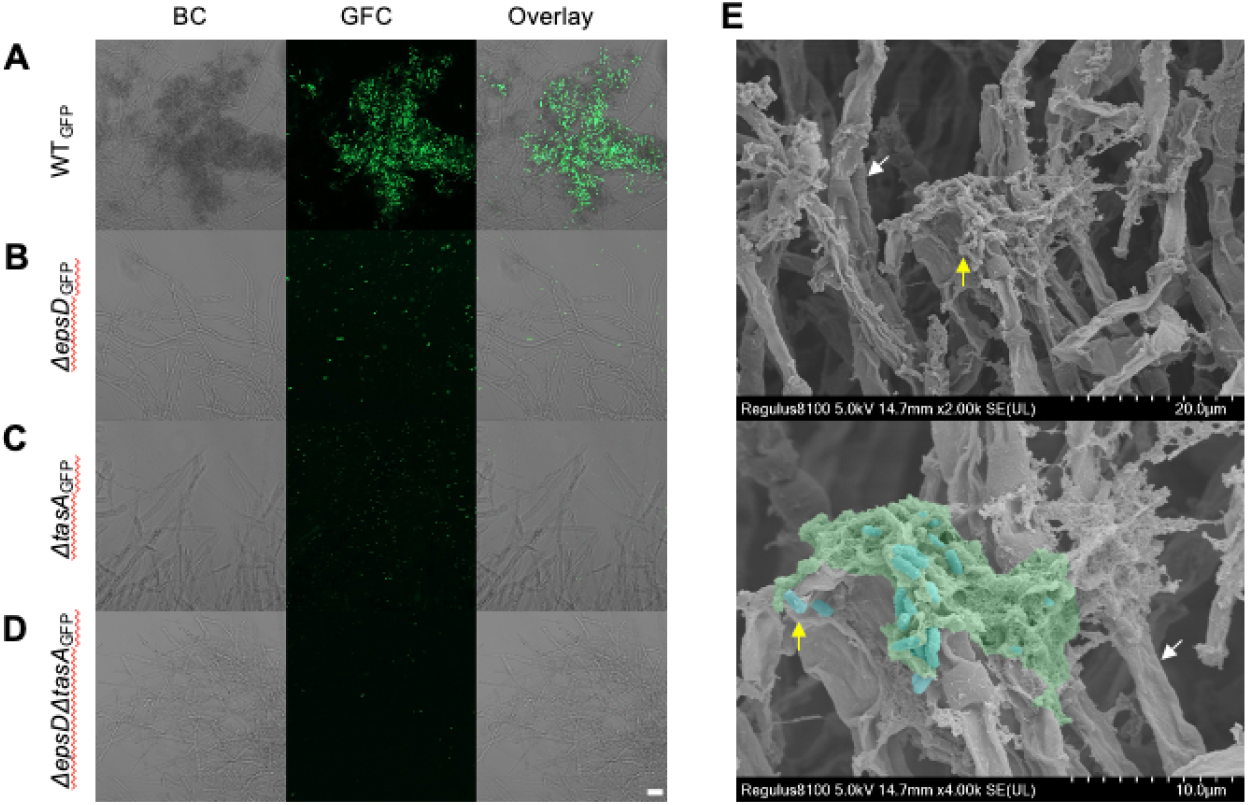
Attachment of *B. velezensis* on *T. guizhouense* hyphae. **(A)** wild type, **(B)** *ΔepsD*, **(C)** *ΔtasA*, **(D)** *ΔepsDΔtasA*, images made by CLSM. BF: bright channel; GFC: green fluorescent channel; RFC: red fluorescent channel. Scale bars indicate 20 μm. **(E)** SEM images of symbiont. White arow: *T. guizhouense*. Yellow arrow and blue color: *B. velezensis*. Green color: EPS of biofilm. Scale bars were showed in figures.

Following this, we conducted a differential gene expression analysis to reveal the significant differences in *B. velezensis* transcriptome during fungal hyphae colonization (Fig S2). Specifically, 1390 genes were upregulated while 1376 genes were downregulated in the *B. velezensis* cells colonizing the hyphae compared with the monoculture cultures (Fig 3D). Conversely, there were only 581 and 576 genes were up- and downregulated in the cocultured free living *B. velezensis* compared with the monocultures (Fig S4). Notably, the most upregulated gene in hyphae-colonized cells compared with the monoculture was the *tasA* gene in *B. velezensis*, which encodes the amyloidogenic protein of the biofilm matrix [34]. Subsequently, the expression of biofilm-related genes of *B. velezensis* [13] was examined in the transcriptome, including the *eps* operon, *bslA, tapA* operon (that contains also the *sipW* and *tasA* genes) (Fig 3D). The *epsA-O* operon encodes the synthesis of exopolysaccharide (EPS) [35], *tapA* and *sipW* encodes the chaperon and signal peptidase of TasA protein required for amyloid fibre assembly [36], while *bslA* encodes a hydrophobin-like protein [37]. Upregulation of all these biofilm-related genes were observed in the hyphae attached cells that suggests enhanced biofilm matrix production in *B. velezensis* during colonization of the fungal hyphae. Additionally, down regulation of *cwlC* expression indicated, which encodes a cell wall hydrolase [38], is consistent with long chain formation during the onset of biofilm development.

In conclusion, transcriptome analysis uncovered the bacterial differentiation by *B. velezensis* in cocultures during hyphae colonization, suggesting that interaction with the hyphae of *T. guizhouense* requires the biofilm matrix components of *B. velezensis*.

### 3.4 Hyphae colonization by *B. velezensis* requires the major matrix components To further investigate the role of biofilm matrix production-related genes during

hyphae colonization, we removed either one or both genes required for the two major matrix components in the GFP-labeled *B. velezensi*s strain. The **Δ***epsD*, **Δ***tasA*, and **Δ***espD***Δ***tasA* mutants were then co-cultured with *T. guizhouense* fungal pellets comparable to above in (Fig 2A). Confocal laser scanning microscopy (CLSM) analysis revealed that the wild-type (WT) *B. velezensis* successfully colonized the hyphae of *T. guizhouense*, whereas none of the mutants were able to colonize the hyphae (Fig 5A & Fig S5). Additionally, scanning electron microscopy (SEM) imaging of the WT-colonized fungal pellet revealed the presence of extracellular polysaccharides and *B. velezensis* cells colonizing the hyphae (Fig 5B).

## 4 Discussion

BFIs have a profound impact on the nutrition and health of host organisms, as both plants and animals harbor a variety of bacterial and fungal communities [39]. A study on onion tissue bioavailability demonstrated that when inoculated with the mycorrhizal fungus *Golmus intaradices, B. subtilis* helper strain significantly increased biomass due to the bacterium’s phosphate-solubilizing abilities, leading to nitrogen and phosphorus accumulation in the plant [40].

*T. guizhouense* NJAU 4742 and *B. velezensis* SQR9 have been extensively studied as both have demonstrated positive influence on plant growth [10]. Therefore, a subsequent inoculations of the two PGPM were explored to see the influence of potential BFIs on growth of tomato plants. Importantly, the order of inoculation had no influence on the positive impact of these PGPM on plant yield in soil. While inhibition was observed in the conventional laboratory assays with the two microorganisms inoculated at the same time at distant locations on an agar medium, such competitive interaction was seemingly absent during inoculation of these two strains to the soil, potentially due to the spatial structure created by the soil particles [41] and the plant roots, or due to distinct nutrient availability that lessens the growth rate of these two microorganisms, allowing a more subtle contact between them.

It is difficult to truly replicate the conditions present in a complex soil environment using a laboratory liquid medium. To circumvent the lack of a reliable system, we customized a coculture method [28,29], in which *T. guizhouense* spores was first allowed to germinate and grow before *B. velezensis* was inoculated. This method provided a stable system to study BFIs. Interestingly, our results indicated *B. velezensis* could colonize on *T. guizhouense* hyphae, contrary to the antagonistic behavior observed when the two microorganisms were inoculated in a distance on PDA medium. Using the coculture system, transcriptome analysis revealed significant differences in gene expression between *B. velezensis* cells colonizing the hyphae versus free living state in the presence or absence of the fungus. Secondary metabolite gene expression was increased in free-living cocultures of *B. velezensis*, suggesting that *B. velezensis* competed for resources, space, and nutrients with *T. guizhouense* by upregulating secondary metabolites [33]. The fungal hyphae colonizing *B. velezensis* cells potentially occupy a distinct ecological niche, due to their reduced mobility and chemotaxis.

As transcriptome analysis of hyphae colonizing bacterial cells suggested an important role biofilm production by *B. velezensis*, mutant strains lacking matrix production were tested and visualize during coculturing using different microscopy methods. As none of the bacterial mutants that lacked the biofilm matrix was able to colonize the fungal hyphae, these experiments clearly demonstrated the essentiality of biofilm matrix production for establishment of BFI under those conditions. The importance of biofilm components in BFIs has been highlighted in previous studies. For example, the biofilm matrix of *B. subtilis* is essential for the colonization of the hyphae of the Ascomycota *Aspergillus niger* and the Basidiomycota *Agaricus bisporus* [29].

The outcome of the interaction between bacteria and fungi depends on natural physical interaction [39]. Bacteria coexist within hyphae in various forms, including free-living cells, hyphae attached single cells, endohyphal cells, and matrix embedded cells of biofilms [42]. Previous studies highlighted the role of biofilm matrix during colonization of the hyphae of *Ascomycetes, Basidiomycetes* and *Zygomycetes* [43]. However, most of the BFI-related biofilm research focused on mycorrhizal fungi, which inherently cooperate with bacteria. For example, intricate interaction between *B. velenzesis* and the arbuscular mycorrhizal fungi, *Rhizophagus irregularis* suggested that the biosynthesis of specific lipopeptides and antimicrobial compounds in *B. velezensis* is attenuated as a mechanism to ensure stable coexistence of these two microoorganisms [44].

Future experiments could explore whether in addition to biofilm-mediated colonization, *B. velezensis* can potentially also exploit the *T. guizhouense* hyphae as a “fungal highway”, similar to those observed between *Aspergillus nidulans* and *B. subtilis* [45] or if there is any growth stimulating metabolites are production by *B. velezensis* and *T. guizhouense*. Concurrently, we speculated that within soil ecosystems, a broader spectrum of bacteria interact with hyphae. Identifying and investigating these beneficial interactions among there strains could advance the theoretical understanding of BFIs.

## 5 Conclusion

The interactions within the soil rhizosphere microbiome are complex, yet they are often underestimated by researchers. Factors, such as the ecological niche differentiation and variations in growth rates are frequently overlooked. Notably, *Bacillus* and *Trichoderma* exhibit distinct interactions in the laboratory while simultaneously promoting plant growth. Here, we propose that mitigating the differences in growth rates could offer a simple solution studying BFI in the laboratory. Our findings elucidated that *B. velezensis* could colonize the hyphae of *T. guizhouense* via biofilm formation with a distinguished transcriptome from non-colonizing bacterial cells. This study provides a novel perspective on BFIs in soil and furnishes a theoretical framework for exploring specific interactions among bacteria, fungi and plants. Such insights are essential for advancing our comprehension of harnessing soil microbiota to improve plant growth and enhance agricultural productivity.

## Data availability

All data are presented within the article.

## Acknowledgements

This work was financially supported by National Nature Science Foundation of China (42090060 and 42307173), and the National Key Research and Development Program (2022YFD1500202, 2022YFF1001800). J.X. was supported by a Chinese Scholarship Council fellowship (202206850025) and Postgraduate Research & Practice Innovation Program of Jiangsu Province (KYCX23_0802). Á.T.K. was supported by a start-up fund from Institute of Biology Leiden and the Novo Nordisk Foundation within the INTERACT project of the Collaborative Crop Resiliency Program (NNF19SA0059360).

