## supplementary figures for "Bridging the Gap: biofilm-mediated establishment of *Bacillus velezensis* on *Trichoderma guizhouense* mycelia"

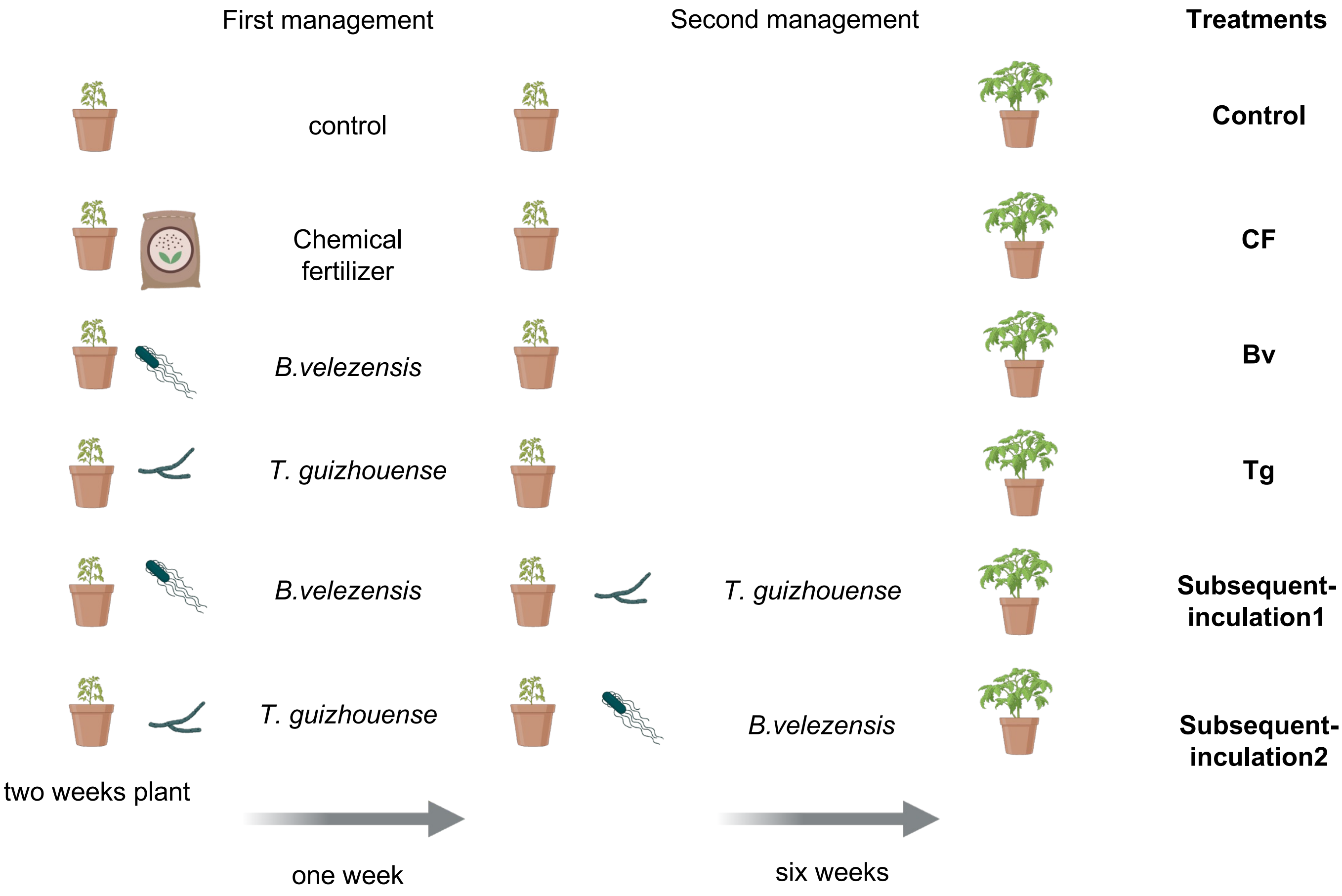


**Fig S1 The tomato plant experiments design*.*** Control: water; CF: chemical fertilizer; Bv: *B. velezensis*; Tg: *T. guizhouense*; Subsequent-inculation1): inoculated *B. velezensis* in the soil first then *T. guizhouense*; Subsequent-inculation2(: inoculated *T. guizhouense* in the soil first then *B. velezensis*.

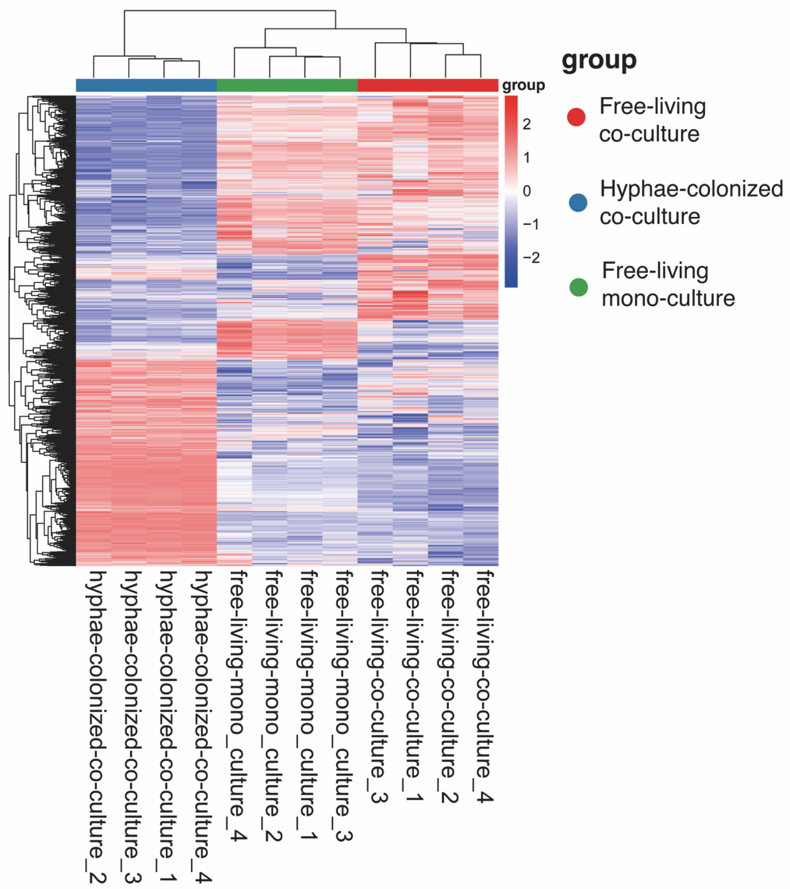


**Fig S2 The differential gene expression analysis of *B. velezensis* in different treatments.**

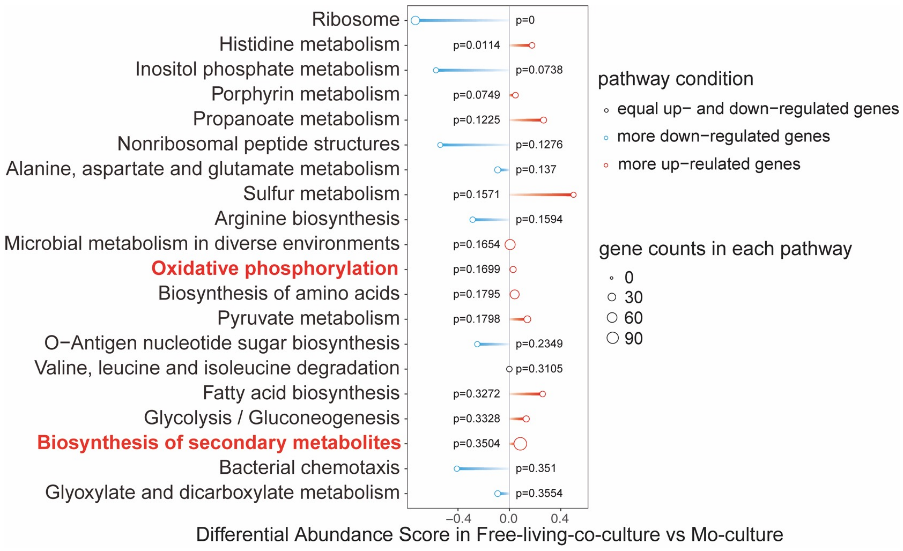


**Fig S3 KEGG enriched pathway analysis of free-living co-cultured *B. velezensis* compared with mono-culture(control).** Blue indicated there were more down-regulated genes in the pathway, red indicated there were more up-regulated genes in the pathway, black indicated equal up and down-regulated genes in the pathway.

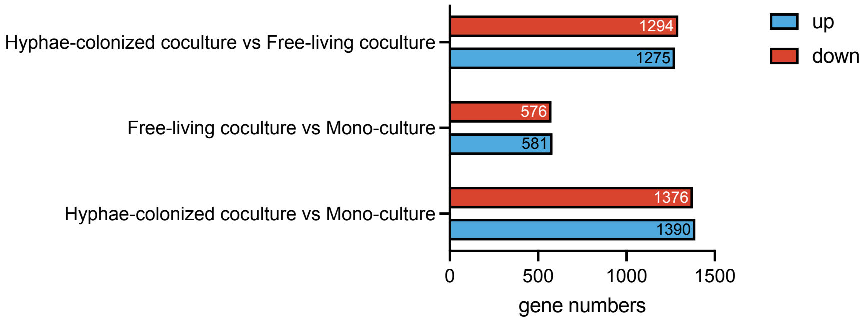


**Fig S4 Numbers of the differentially regulated genes in three treatments**. LFC >2, $p$ <0.05, t test.

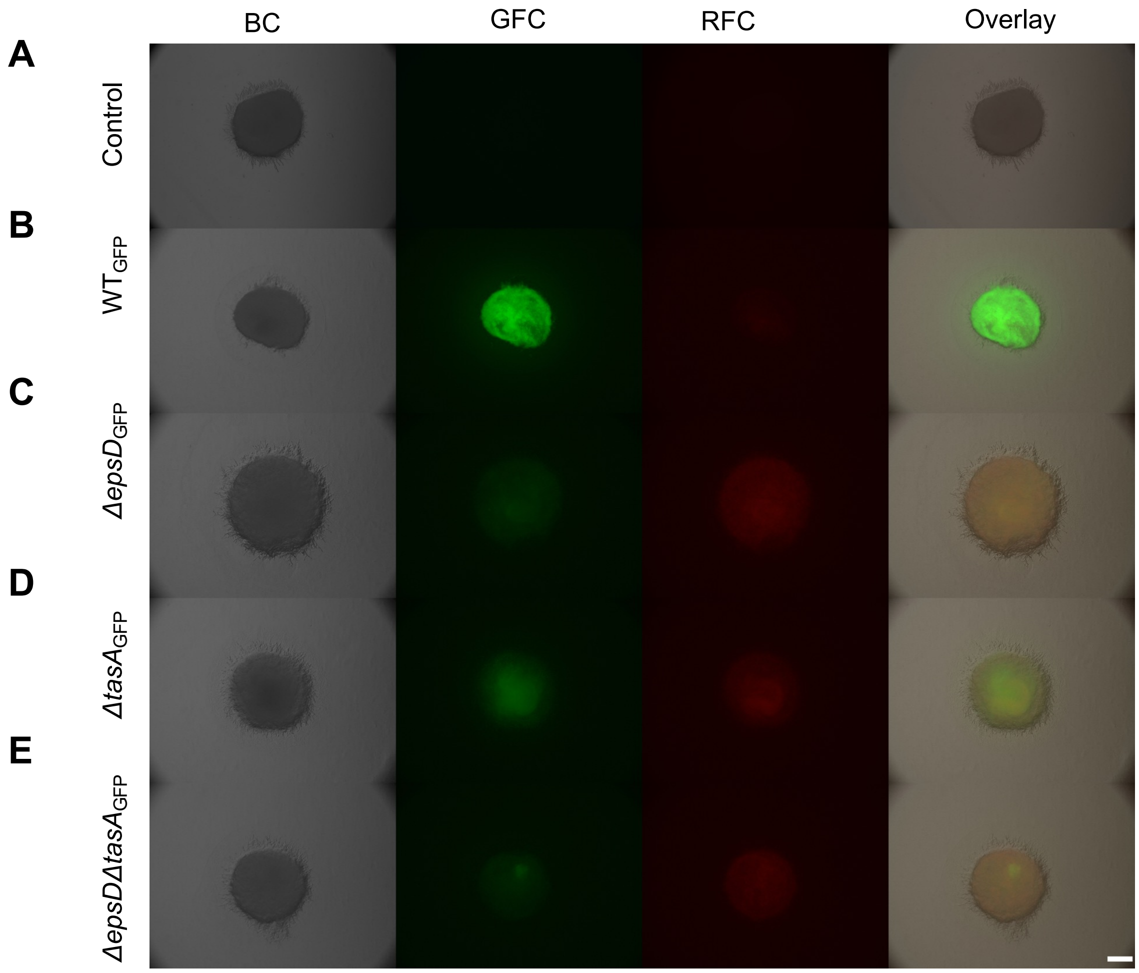


**Fig S5 Attachment of *B. velezensis* on *T. guizhouense* hyphae. (A)**control, **(B)** wild type, **(C)** *ΔepsD*, **(D)** *ΔtasA*, **(E)** *ΔepsDΔtasA,* images made by Stereo Microscopy. BC: bright channel; GFC: green fluorescent channel; RFC: red fluorescent channel. Scale bars indicate 500 μm.

### Table S1: Primers used in this study

| Primers | Sequence(From 5' to 3') | Experimental purpose |
|---|---|---|
| tasA_UF | TCGCATGCGATGAATCACTTC | To amplify the upstream region of *tasA* gene in *B. velezensis* |
| tasA_UR | ACTTTTCGGGGAAATGTGCAAGTTCCAAGGCAATGCGA | To amplify the upstream region of *tasA* gene in *B. velezensis* |
| epsD_UF | CGTCTGCATGCCTGAAAAACG | To amplify the upstream region of *epsD* gene in *B. velezensis* |
| eps_UR | CGTTACGTTATTAGTTATGAAAAGCTGTACCGCTCCCC | To amplify the upstream region of *epsD* gene in *B. velezensis* |
| tasA_DF | TATTTAACGGGAGGAAATAATCATTAAATGCGGCCCATGT | To amplify the down stream region of *tasA* gene in *B. velezensis* |
| tasA_DR | ATCGCTTCTCAGCAGCAGC | To amplify the down stream region of *tasA* gene in *B. velezensis* |
| epsD_DF | TATAGCATACATTATACGAGATTCATATTGCCGCCTGC | To amplify the downstream region of *epsD* gene in *B. velezensis* |
| epsD_DR | CATACCGAGCTGTCCGCA | To amplify the downstream region of *epsD* gene in *B. velezensis* |
| Em_F | TCGCATTGCCTTGGAACTTGCACATTTCCCCGAAAAGT | cloning |
| Em_R | ACATGGGCCGCATTTAATGATTATTTCCTCCCGTTAAATA | cloning |
| Spc_F | GGGGAGCGGTACAGCTTTTCATAACTAATAACGTAACG | cloning |
| Spc_R | GCAGGCGGCAATATGAATCTCGTATAATGTATGCTATA | cloning |
